# Conflicts between the DNA replication and repair machineries promote cell death in Gram-positive bacteria

**DOI:** 10.1101/2024.10.24.619994

**Authors:** Hannah Gaimster, Daniel Stevens, James Grimshaw, Julia Hubbard, Katarzyna Mickiewicz, Heath Murray, Charles Winterhalter

## Abstract

Cellular proliferation relies on successful coordination and completion of genome replication and segregation. To help achieve this, many bacteria utilise regulatory pathways that ensure DNA replication initiation only occurs once per cell cycle. When dysregulated, loss of DNA replication control can have severe consequences. In *Escherichia coli* it has been established that hyper-initiation of DNA synthesis leads to pleiotropic genome instability and cell death. Therefore, targeting DNA replication initiation proteins to promote hyper-initiation may be an approach to generate novel antimicrobials. However, the pathways and potential consequences of replication hyper-initiation in Gram-positive species remain enigmatic. To address this question, we devised genetic systems to artificially induce hyper-initiation in the model organism *Bacillus subtilis* and the pathogen *Staphylococcus aureus*. In both species, hyper-initiation elicited cellular degeneration culminating in growth inhibition by cell death. During this process in *B. subtilis*, temporal analyses revealed the early onset of the DNA damage response, followed by membrane depolarisation and cell lysis. This phenotype could be supressed by removing pathways that repair damaged DNA, suggesting that cell death is a consequence of conflicts between DNA replication and repair. In *S. aureus,* cells quickly accumulated striking morphological changes associated with rapid loss of chromosomal DNA and death via a lysis-independent pathway. Moreover, inducing hyper-initiation in *S. aureus* was observed to decrease bacterial survival during infection of murine macrophages. Taken together, the data suggest that stimulating initiation of bacterial DNA synthesis could be an alternative approach to inhibiting microbial growth, particularly in combination with compounds that inhibit or poison DNA repair, akin to cancer therapies.

## INTRODUCTION

A successful cell cycle requires replication and segregation of the genome. In most cells, DNA synthesis begins at specific chromosomal loci called origins and requires accurate coordination with chromosome segregation and cell division to produce viable daughter cells [1, 2]. Importantly, while replication initiation culminates in loading of ring-shaped helicases enabling double-strand DNA unwinding prior to synthesis, the molecular mechanisms required to achieve helicase loading in prokaryotes and eukaryotes are distinct [3]. Therefore, the unique and essential characteristics of the bacterial DNA replication initiation pathway make it an attractive target for drug development [4].

In bacteria, chromosome replication generally initiates from a single origin (*oriC*), where the universal prokaryotic initiator protein DnaA binds to sequence specific elements (DnaA-boxes and DnaA-trios) and unwinds the DNA duplex [5–7], followed by recruitment and loading of helicases around single-stranded DNA [8–10]. During this process, DnaA activity is tightly regulated to ensure genome replication occurs only once per cell cycle, while bidirectional DNA synthesis proceeds up to a region called the terminus where converging replication forks collide [11–13]. Failure to maintain this homeostasis can lead to dramatic consequences affecting a range of cellular processes resulting in genomic instability [14, 15].

Abnormal initiation of genome duplication can result in either replication inhibition or excessive DNA synthesis, both of which can inhibit bacterial proliferation. Intuitively, under-initiation of DNA replication produces cells that lack a copy of the genome [16]. A more complex situation is observed during hyper-initiation of DNA replication. Excessive genome replication has been extensively studied and remains an active field of research in the Gram-negative model organism *Escherichia coli,* where under aerobic growth conditions hyper-initiation causes DNA double-strand breaks (DSBs) as a result of the DNA replication machinery (replisome) encountering repair systems acting on the genome [17, 18]. These detrimental events affect the normal cell cycle and can result in significant cell death via improper completion of DNA synthesis or chromosome segregation defects, yet *E. coli* may employ several strategies to overcome lethal over-initiation [19–22]. However, the lethal phenotype associated with hyper-initiation of DNA replication remains enigmatic, as suppression of cell death did not correlate with a significant decrease in DSBs [17]. Moreover, it is still unclear whether hyper-initiation of DNA replication generally inhibits bacterial proliferation in Gram-positive bacteria, and if so, the specific pathways leading to potential cell death are yet to be determined across species.

In this study, we investigated the consequence of DNA replication hyper-initiation in the Gram-positive model organism *Bacillus subtilis* and the opportunistic pathogen *Staphylococcus aureus.* In both cases hyper-initiation resulted in loss of cell viability, however the lethal phenotypes were not identical. The *B. subtilis* strain displayed cell elongation, DNA damage, membrane depolarisation, and eventually cell lysis. The *S. aureus* strain also displayed altered cell morphology with clear defects in chromosome inheritance, but here loss of cell viability occurred through a lysis-independent mechanism. Interestingly, induction of *S. aureus* hyper-initiation inhibited bacterial growth in a murine macrophage model, highlighting the potential of stimulating DNA replication initiation as an approach to reduce bacterial load during infection.

## MATERIALS AND METHODS

### Reagents and growth conditions

Nutrient agar (NA; Oxoid) was used for routine selection and maintenance of *E. coli* and *B. subtilis* strains. Tryptic soy agar (TSA; Oxoid) was used for routine selection and maintenance of *S. aureus* strains. Unless otherwise stated, *B. subtilis* and *S. aureus* strains were grown at 30°C and *E. coli* was grown at 37°C. Supplements were added as required: ampicillin (100 µg/ml), erythromycin (1 µg/ml) in conjunction with lincomycin (25 μg/ml), chloramphenicol (5 µg/ml for *B. subtilis*, 10 µg/ml for *S. aureus*), kanamycin (5 µg/ml), spectinomycin (50 µg/ml), tetracycline (10 μg/ml), xylose (1% w/v) and anydrotetracycline (aTc, 20 ng/ml). All chemicals and reagents were obtained from Sigma-Aldrich. RAW-Blue cells were maintained in Dulbecco’s Modified Eagle’s medium (DMEM, Sigma Cat#D6429) supplemented with 5% fetal bovine serum (FBS) at 37°C and 5% CO_2_.

### Biological resources: *E. coli*, *B. subtilis* and *S. aureus* strains

All strains and cell lines used in this study are listed in Supplementary Table 1.

### *B. subtilis* strain construction

Transformation of competent *B. subtilis* cells was performed using an optimized two-step starvation procedure as previously described [23]. Briefly, recipient strains were grown overnight at 30°C in transformation medium (Spizizen salts supplemented with 1 μg/ml Fe-NH_4_-citrate, 6 mM MgSO_4_, 0.5% w/v glucose, 0.02 mg/ml tryptophan and 0.02% w/v casein hydrolysate) supplemented with xylose where required. Overnight cultures were diluted 1:20 into fresh transformation medium supplemented with xylose where required and grown at 30°C for 3 hours with continual shaking. An equal volume of prewarmed starvation medium (Spizizen salts supplemented with 6 mM MgSO_4_ and 0.5% w/v glucose) was added and the culture was incubated at 30°C for 2 hours with continuous shaking. DNA was added to 350 μl cells and the mixture was incubated at 30°C for 1 hour with continual shaking. 20-200 μl of each transformation was plated onto selective media supplemented with xylose where required and incubated at 30°C for 24-48 hours. Note that strains harbouring the *dnaA^G154S^*/*parA^G12V^*hyper-initiation system and/or *mutM*/*mutY* knockouts are prone to accumulation of suppressor mutations upon storage or repeated rounds of propagation. For best practice, it is recommended to rebuild these strains using sequence-validated recombinant DNA and to perform whole genome sequencing on candidate colonies prior to use.

### *S. aureus* strain construction

Transformation of competent *S. aureus* cells was performed using plasmid electroporation [24]. Briefly, recipient cells were grown overnight in tryptic soy broth (TSB; Oxoid). Overnight cultures were diluted 1:100 in fresh TSB and grown to an absorbance at 600 nanometers (A_600_) of 0.6. Cells were rapidly cooled down in an ice water bath and washed three times using an equal volume of ice-cold sterile deionised water followed by two washes in an equal volume of ice-cold 10% w/v glycerol solution. After the final centrifugation, cells were resuspended in 1:100 of the initial culture volume in ice-cold 10% w/v glycerol, snap frozen using liquid nitrogen, and stored at -80°C for up to six months. Before electroporation, electrocompetent cells were thawed at room temperature for 5 min, centrifuged at 10,000 g for 1 min, and resuspended in 100 μl of electrocompetent buffer (10% w/v glycerol supplemented with 500 mM sucrose). DNA was added to the cell mixture and electroporation was performed at 2.3 kV, 100□Ω, 25 μF (Bio-rad Gene Pulser II) using a 1 mm gap cuvette (VWR #732-1135). Cells were immediately resuspended in 1 ml of TSB supplemented with 500 mM sucrose, incubated for 1.5 hours at the permissive temperature, plated onto selective medium and incubated at permissive temperature for 24-48 hours.

### *E. coli* plasmid construction and propagation

*E. coli* transformation was performed in DH5α via heat-shock following the Hanahan method [25]. All plasmids were sequenced and descriptions of plasmid construction, where necessary, are provided below.

pDS150 was generated using Gibson assembly to introduce *saDnaA* into pRAB11 [26].

pDS157 was created using Quickchange mutagenesis on pDS150 to introduce the *saDnaA^G159S^* point mutation.

### Spot titre assays

*B. subtilis* cells were grown in penassay broth medium (PAB) overnight at 30°C in the presence of xylose. Overnight cultures were diluted 1:100 into fresh PAB with or without xylose (time = 0 h). Samples were harvested at time points indicated, 10-fold serial dilutions were made into PAB with xylose, and 5 μL aliquots were spotted onto NA plates supplemented with xylose. For *S. aureus* spot titre assays, cells were grown in TSB supplemented with chloramphenicol overnight at 30°C and 5 μL of serial dilutions were spotted onto TSA supplemented with chloramphenicol in the presence or absence of aTc. All plates were incubated at 30°C for 24 hours. Experiments were performed independently at least three times and representative data are shown.

### Automated plate reader analyses

Strains were grown overnight at 30°C in PAB with xylose (*B. subtilis*) or TSB supplemented with chloramphenicol (*S. aureus*). Overnight cultures were diluted 1:100 into PAB with or without xylose (*B. subtilis*), or 1:1000 into TSB with or without aTc (*S. aureus*) and automated absorbance measurements were captured over time at 30°C using a Tecan Sunrise plate reader (high shaking parameters with 2.8 mm shake width at 12.3 Hz, absorbance measurement every six minutes). All experiments were independently performed at least three times and representative data are shown.

### Sample preparation for marker frequency analysis

*B. subtilis* cells were grown overnight in PAB at 30°C with xylose. Overnight cultures were diluted 1:100 into PAB with or without xylose (time = 0 h) and incubated at 30°C. For *S. aureus*, cells were grown overnight in TSB in the presence of chloramphenicol, followed by 1:100 dilution into TSB with or without aTc for two hours before harvesting samples. Samples (500 μl) were harvested at times indicated and immediately mixed with sodium azide (1% w/v final concentration) to arrest growth and genome replication. Cultures were collected by centrifugation, the supernatant discarded, and pellets were flash frozen in liquid nitrogen before genomic DNA (gDNA) extraction using the DNeasy blood and tissue kit (Qiagen).

### Quantitative PCR (qPCR)

qPCR was performed using the Luna qPCR mix (NEB) to measure the relative amount of origin DNA compared to the terminus. All PCR reactions were run in a Rotor-Gene Q instrument (Qiagen) using 20 μl reaction volumes in a Rotor-Disc 100 (Qiagen). Standard curves were obtained using the Rotor-Gene Q Software v2.0.2 (Qiagen) to calculate the efficiency of each primer pair, which varied ∼5% between sets. Oligonucleotide primers designed to amplify *incC* (qSF19/qSF20 for *B. subtilis* or oDS263/oDS264 for *S. aureus*) and the terminus (qPCR57/qPCR58 for *B. subtilis* or oDS269/oDS270 for *S. aureus*) were typically 20-25 bases in length and amplified a ∼100 bp PCR product (Supplementary Table 2). Individual *ori:ter* ratios were obtained in three steps: first, every Ct value was converted to 1/2^Ct^ and technical triplicates were averaged to generate a single enrichment value; second, origin enrichment was normalised by corresponding terminus values; third, relative *ori:ter* values were normalised by the enrichment obtained in control conditions. For *B. subtilis*, data was normalised to spore DNA as non-replicating control (*ori:ter* ratio = 1). For *S. aureus*, data was normalised to cells grown with the pRAB11 plasmid (empty vector used for the construction of *saDnaA* variants) harvested at the same time point in exponential phase. Error bars indicate the standard error of the mean for 2-4 biological replicates.

### Statistical analyses

Statistical analyses were performed using Student’s t-tests and significance of p-values is displayed on individual figure panels and explained in figure legends. The exact value of n is given in method details and represents the number of biological repeats for an experiment. Tests were based on the mean of individual biological replicates. Differences were considered as significant if their associated p-value was below 0.05.

### Microscopy and image analysis

Microscopy was performed on an inverted epifluorescence microscope (Nikon Ti) fitted with a Plan Apochromat Objective (Nikon DM 100x/1.40 Oil Ph3). Light was transmitted from a CoolLED pE-300 lamp through a liquid light guide (Sutter Instruments) and images were collected using a Prime CMOS camera (Photometrics). Fluorescence filter sets were from Chroma: DAPI (49000, EX350/50, DM400lp, EM460/50), GFP (49002, 537 EX470/40, DM495lpxr, EM525/50) and mCherry (49008, EX560/40, 538 DM585lprx, EM630/75). Digital images were acquired using Metamorph (version 7.7) and NIS-Elements software. Cells were mounted on ∼1.2% agar pads and a 0.13-0.17 mm glass coverslip (VWR) was placed on top. Wavelengths used: Brightfield and mCherry (150 ms), GFP/DAPI (50 ms), all at 100% exposure.

For timelapse microscopy experiments, a GeneFrame was used to create a flat agarose pad using 0.7% agarose dissolved into PAB with or without xylose (*B. subtilis*) or TSB with or without aTc (*S. aureus*). An aliquot of cells (1.5 μL) was spotted onto the pad and a 0.13-0.17 mm glass coverslip (VWR) was placed on top. Digital images were acquired every 15 minutes for 8-12 hours.

For cultures stained with DiBAC_4_(3) (Invitrogen, Thermofisher), 0.5 μl dye was added to 100 μl of cells and the mixture was incubated at 30°C with shaking at 800 rpm for 5 minutes using a Thermomixer C (Eppendorf), then spotted onto an agarose slide. DiBAC_4_(3) preferentially binds cellular membranes that are depolarised, emitting a green fluorescence signal to allow for detection (i.e. cells appear fluorescent if the cell envelope is compromised).

For cultures stained with 4′,6-diamidino-2-phenylindole (DAPI, Thermofisher), the dye was added to 100 μl of cells at a final concentration of 1 μg/ml and the mixture was incubated at 30°C with shaking at 800 rpm for 5 minutes using a Thermomixer C (Eppendorf), then spotted onto an agarose slide.

Fiji software was used for initial image analysis [27]. Brightness and contrast used for each time point and condition remained consistent and all images are scaled to 0.065 microns. Cell segmentation was performed on phase-contrast images using Omnipose with the *bact_phase_omni* model, further refined for *S. aureus* using custom training [28]. Image processing and analysis were conducted in scikit-image [29]. Objects smaller than 0.845 μm² (*B. subtilis*) or 0.634 μm² (*S. aureus*) were excluded, and mean background fluorescence was subtracted. Nucleoids were segmented with Ilastik, with labels manually corrected using the membrane channel [30]. Anucleate *B. subtilis* cells were defined as those with nucleoid area below 5% of individual cell sizes.

### Macrophage infection with *S. aureus* and gentamycin protection assay

RAW-Blue cells were seeded the day before bacterial infection experiments at 0.5x10^6^ cells per ml in DMEM 5% FBS medium in 25 cm angled neck flasks (NUNC). Macrophages were challenged at a multiplicity of infection (MOI) of 1:5 in duplicates for *S. aureus* infection with CW1150 or CW1151 bacterial strains. The challenge was carried out in 10 mL of DMEM 5% FBS and cells were incubated for 1 hour at 37°C and 5% CO_2_ to allow bacterial uptake by the macrophages. Following this incubation, gentamycin was added to all flasks (100 ug/ml final concentration) to kill extracellular bacteria and aTc was supplemented to one of each of the duplicates (400 ng/ml final concentration) to induce SaDnaA variant overexpression. Flasks were then incubated for 20 hours at 37°C and 5% CO_2_. Adherent cells were dislodged with a scrapper and 1 ml was transferred into an Eppendorf tube and spun down at 13,000 rpm. The supernatant was aspirated and the pellet was resuspended by pipetting up and down in 1 ml of Triton X100 (0.5% v/v) to lyse macrophages. Serial dilutions were performed and plated on TSA supplemented with 10 ug/ml chloramphenicol for selection of *S. aureus* carrying the inducible plasmid. Bacterial cells were incubated overnight at 37°C and used to quantify CFUs for survival plots. Macrophage experiments were performed as biological duplicates.

## RESULTS

### DNA replication hyper-initiation causes cell lysis in *B. subtilis*

A genetic system was employed to control the frequency of DNA replication initiation in *B. subtilis.* This system exploits a hyperactive variant of DnaA (DnaA^G154S^) that causes cells to over-initiate DNA replication [31]. To control hyper-initiation, a variant of the regulator ParA (ParA^G12V^) was employed to inhibit DnaA activity [32]. The experimental strain encodes *dnaA^G154S^*at the endogenous locus (under the control of its native expression system) and *parA^G12V^* at an ectopic locus (under the control of a xylose-inducible promoter). Hereafter, we will refer to this strain (*dnaA^G154S^ P_xyl_-parA^G12V^*) as *Bs^HI^* (for *B. subtilis* hyper-initiation). The *Bs^HI^* strain was cultured in the presence of xylose to repress hyper-initiation, followed by growth in the absence of xylose to repress and dilute ParA^G12V^, thereby enabling full activity of the hypermorphic DnaA^G154S^ variant (Fig. 1A).

**FIGURE 1:**
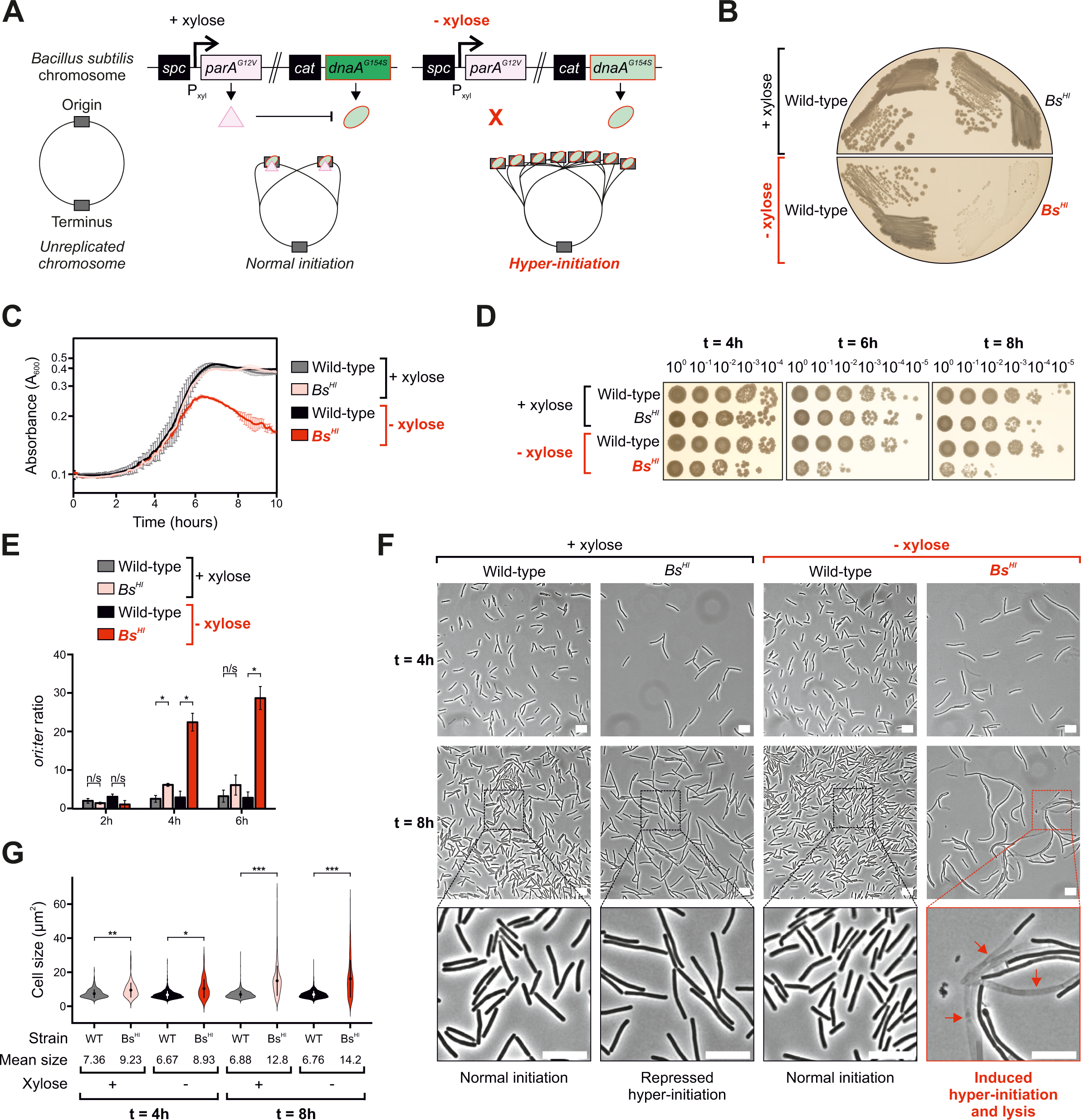
Hyper-initiation of DNA replication causes lysis and cell death in *B. subtilis*. **(A)** Genetic system used to artificially modulate DNA replication hyper-initiation. Xylose induction controls expression of *parA^G12V^*, which enables ParA^G12V^ to downregulate the activity of the hyperactive DnaA^G154S^ variant resulting in repression of hyper-initiation. In the absence of xylose, *parA^G12V^*is no longer expressed and DnaA^G154S^ allows *B. subtilis* to hyper-initiate. **(B)** Restreaks showing the phenotype associated with *dnaA^G154S^ P_xyl_-parA^G12V^* (*Bs^HI^*) cells in the absence of xylose after 24 h growth. **(C)** Growth curves showing the arrest of cell growth in *Bs^HI^*cells after 6 h incubation in the absence of xylose. **(D)** Spot titre analyses showing the loss of viability observed in *Bs^HI^* cells after 6 h growth in the absence of xylose. **(E)** Marker frequency analyses demonstrate that *Bs^HI^* cells start to hyper-initiate DNA replication after 4 h growth in the absence of xylose. **(F)** Phase-contrast microscopy showing the lysis phenotype observed in *Bs^HI^* cells after 8 h growth in the absence of xylose. Scale bar indicates 10 μm. Red arrows highlight lysed cells. **(G)** Microscopy analyses showing quantification of individual cell sizes from the data shown in (F). Each condition highlights mean cell size (circles within violins) and standard deviation (vertical line crossing the mean). (B-G) Strains: Wild-type/WT (168CA), *Bs^HI^* (HM946). Non-significant (n/s), P values: < 0.05 (*), < 0.001 (**), < 0.0001 (***).

Using this system, strains were streaked onto solid growth medium with or without xylose and incubated overnight. While both wild-type and *Bs^HI^* strains showed similar growth in the presence of inducer, the *Bs^HI^*strain displayed a severe growth defect in the absence of xylose (Fig. 1B). To further characterise this phenotype, cell growth in liquid media was followed over time. The results confirmed that the *Bs^HI^* strain had a similar growth rate to wild-type in the presence of xylose, whereas *Bs^HI^* cells could not reach the same density and appeared to lyse after ∼6 hours in the absence of the inducer (Fig. 1C).

The impact of hyper-initiation on cell viability was quantified by growing cells in liquid medium with or without xylose, followed by aliquoting a serial dilution of each sample onto growth medium containing xylose (i.e. to repress hyper-initiation and allow growth of any viable cells). This spot titre analysis showed a similar number of colony forming units (CFU) between wild-type and *Bs^HI^*strains after four hours of growth. However, by six and eight hours of growth without xylose, the *Bs^HI^* strain displayed a >100-fold decrease in CFU compared to both wild-type and *Bs^HI^* cultured in the presence of xylose (Fig. 1D).

Marker frequency analysis (MFA) was used to determine the levels of DNA replication initiation under these experimental conditions. Over a six-hour time course, MFA showed *ori:ter* ratios ranging from 2.0 to 3.3 for wild-type *B. subtilis* (Fig. 1E). During lag phase, the *Bs^HI^*strain showed comparable *ori:ter* ratios to wild-type regardless of the presence or the absence of inducer (two-hour timepoint, Fig. 1E). Strikingly, *Bs^HI^* cells displayed a significantly increased *ori:ter* ratio following four hours of growth without xylose (*ori:ter* = 22 in exponential phase), whereas the same strain grown in the presence of inducer only mildly over-initiated (*ori:ter* = 6 compared to wild-type *ori:ter* = 2.5, Fig. 1E). The results contrast with a previous report of hyper-initiation in *B. subtilis* where cells with a milder increase in replication initiation remained viable [33], which indicates that the hyperactive *dnaA^G154S^* allele is more penetrant. Taken together, the data suggest that under this growth regime of depleting ParA^G12V^ to allow full activity of DnaA^G154S^, hyper-initiation of DNA replication precedes significant cell death.

Phase contrast microscopy was used to investigate the fate of single *B. subtilis* cells experiencing DNA replication hyper-initiation. Regardless of the presence or absence of xylose in conditions preceding significant cell death (e.g. four-hour time point), *Bs^HI^*cells showed a ∼25% increase in cell size compared to wild-type (Fig. 1F-G), indicating that the normal cell cycle is perturbed in this strain [34]. We attribute morphological defects observed in the presence of inducer to pleiotropic activities of ParA^G12V^, which is known to affect other cellular processes (summarised in [35]). Consistent with inherent stress under these growth conditions, *Bs^HI^* cell size was further increased to approximately twice the size of wild-type by eight hours of growth (Fig. 1G). However, incubation in the absence of xylose resulted in dramatic changes to the cell envelope (“bulging” phenotype) with many cells appearing phase light, a hallmark of cell lysis (Fig. 1F). These phenotypes can be seen in greater detail in time lapse videos of *Bs^HI^*cells grown in the presence or absence of xylose (Supplementary Videos 1 and 2, respectively). Together, these data indicate that in *B. subtilis,* DNA replication hyper-initiation causes a loss of viability via cell lysis.

### Hyper-initiation activates the DNA damage response and leads to membrane depolarisation

In bacteria it is established that severe DNA damage elicits a stress response (the SOS response), allowing time to repair the damage and segregate replicating chromosomes [34, 36, 37]. To induce the SOS response, the critical recombination protein RecA binds to single-stranded DNA and stimulates the autocleavage of LexA, the transcriptional repressor of the SOS regulon [38]. Upon cleavage, LexA can no longer bind to DNA to repress transcription, resulting in induction of the SOS response.

One of the SOS responsive genes most strongly upregulated in *B. subtilis* encodes the cell division inhibitor YneA [39]. We hypothesised that the cell size increase and lysis observed during hyper-initiation (Fig. 1F) may be caused by induction of the DNA damage response. To test this model, a fluorescent transcriptional reporter under control of the LexA-dependent *yneA* promoter (*P_yneA_-gfp*) was transformed into *Bs^HI^* (Fig. S1A). As controls, it was confirmed using spot titre assays that addition of the *yneA* reporter cassette did not alter the phenotype associated with hyper-initiation (Fig. S1B) and that YneA was not causative of hyper-initiation induced cell death (Fig. S2A-B). Following growth in liquid media with or without xylose throughout the time window separating the onset of hyper-initiation from accumulation of growth defects leading to cell death (Fig. 1E-G, four to eight hours), expression of *P_yneA_-gfp* was observed using fluorescence microscopy. It was validated that induction of the DNA damage response is a rare event in wild-type cells in the presence or the absence of xylose (Fig. S3A-B). While a minority of *Bs^HI^* cells grown in the presence of xylose appeared to express the GFP reporter, in sharp contrast the majority of cells displayed a GFP signal gradually increasing following depletion of ParA^G12V^ from four hours onwards (Figs. 2A-B, S3C). Induction of the SOS response suggests that DNA replication hyper-initiation leads to the production of DNA damage intermediates (i.e. single-stranded DNA) recognised by RecA.

**FIGURE 2:**
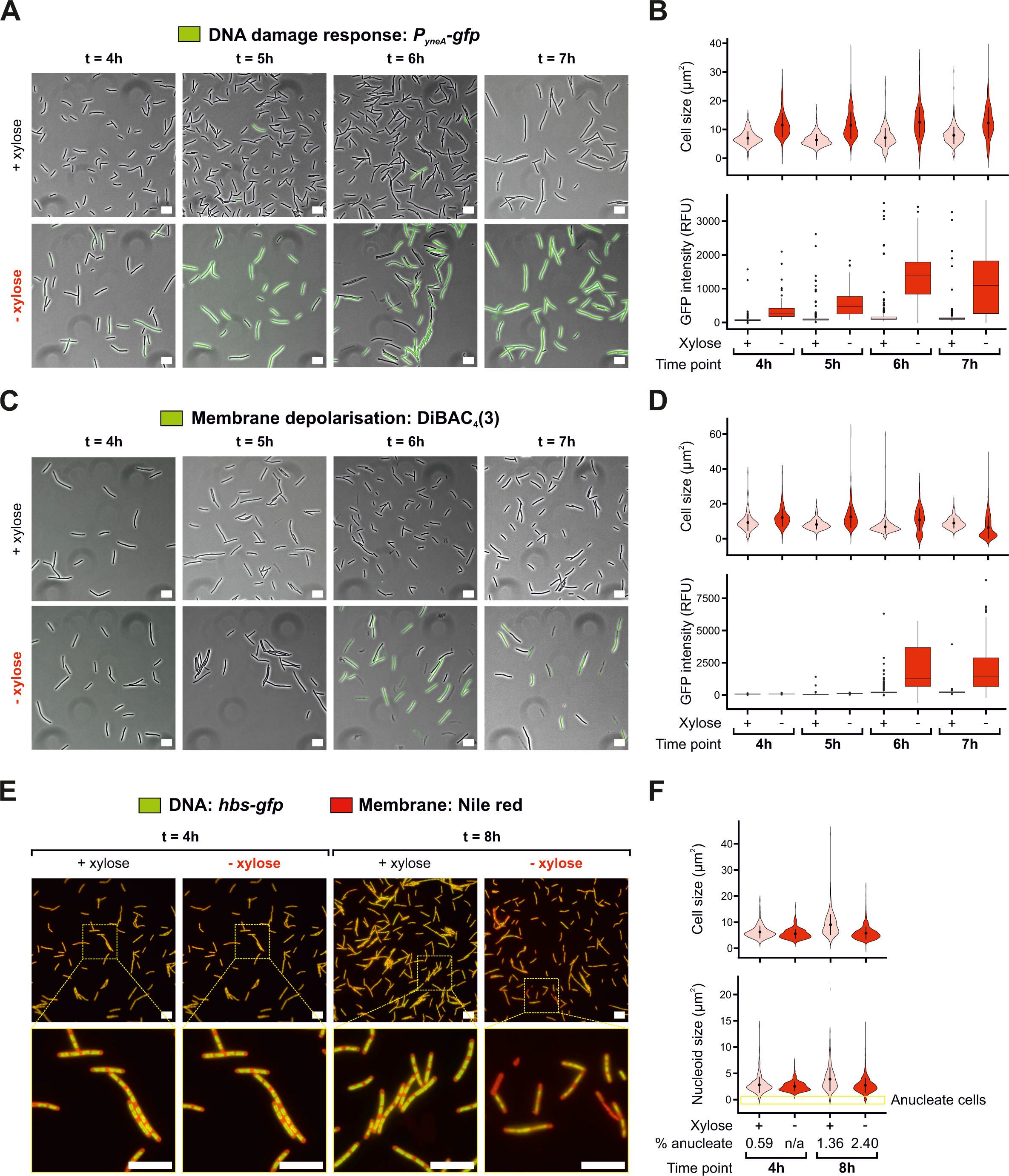
Hyper-initiation causes DNA damage followed by membrane depolarisation and cell lysis via an apoptotic-like mechanism. **(A)** Microscopy experiments showing activation of the DNA damage response in *Bs^HI^* cells in the presence/absence of xylose to control hyper-initiation. Phase contrast images are overlayed with fluorescence signal corresponding to expression of the *P_yneA_-gfp* SOS regulon reporter. Strain: HG287. **(B)** Microscopy analyses showing quantification of individual cell sizes (top) and GFP fluorescence (bottom, DNA damage reporter signal) from the data shown in (A). **(C)** Microscopy experiments showing membrane depolarisation in *Bs^HI^* cells in the presence/absence of xylose to control hyper-initiation. Phase contrast images are overlayed with fluorescence signal corresponding to DiBAC_4_(3) entering depolarised cells. Strain: HM946. **(D)** Microscopy analyses showing quantification of individual cell sizes (top) and GFP fluorescence (bottom, membrane depolarisation signal) from the data shown in (C). **(E)** Microscopy experiments showing the nucleoid in *Bs^HI^* cells in the presence/absence of xylose to control hyper-initiation. Images are merged between green fluorescence for visualisation of DNA content (Hbs-GFP signal) and red fluorescence for membrane stain (Nile red dye). Strain: HM1974. **(F)** Microscopy analyses showing quantification of individual cell sizes (top) and areas covered by the nucleoid (bottom, highlights the number of anucleate cells) from the data shown in (E). n/a indicates that no anucleate cells were found. (A/C/E) Scale bar indicates 10 μm. (B/D/F) Each condition highlights mean cell size (circles within violins), standard deviation (vertical line crossing the mean) and boxplots indicate median lines (within boxes) and outliers (circles outside boxes).

It is established that RecA plays a role during hyper-initiation in *E. coli* but whether this function is conserved in Gram-positive species had not been fully elucidated [18, 40]. To test the importance of RecA during hyper-initiation in *B. subtilis*, we constructed a complementation system by placing an ectopic copy of *recA* under the control of a constitutive promoter (*P_veg_-recA*). It was confirmed that introducing this cassette in *Bs^HI^* cells yielded a similar number of CFU to control strains in the presence or the absence of xylose (Fig. S4A). Spot-titre analyses revealed that deletion of *recA* resulted in synthetic lethality in *Bs^HI^* cells grown in the absence of xylose whereas the presence of *P_veg_-recA* was able to rescue this phenotype (Fig. S4B), indicating that RecA is causative of growth sensitisation under these conditions. Consistent with *E. coli* literature [18], the results suggest that RecA is essential during hyper-initiation in *B. subtilis* to prevent replication fork collapse.

Under conditions enabling hyper-initiation, time course and timelapse microscopy indicated that morphological abnormalities and lysis occur after DNA damage is generated (Fig. 1F and Supplementary Video 2). To further explore the observed compromise to cell envelope integrity, strains were incubated with the fluorescent dye DiBAC_4_(3), which preferentially enters cells with depolarised phospholipid membranes [41]. Under the conditions tested to investigate hyper-initiation in liquid medium, fluorescence microscopy indicated that membrane depolarisation is a rare event in wild-type *B. subtilis* (Fig. S5A-B). However, a significant number of *Bs^HI^* cells displayed fluorescent signals following depletion of ParA^G12V^ for six hours (Fig. 2C-D). Note that a size reduction was observed in *Bs^HI^* cells grown in the absence of xylose for samples matching the onset time of lysis and onwards (Figs. 2D, S5C), which can be attributed to uneven DiBAC_4_(3) staining further affecting morphology and phase detection of compromised cells [41]. Using an *hbs-gfp* cassette to visualise DNA [42], after eight hours of growth in the absence of xylose the *Bs^HI^* strain showed abnormal chromosome content and a significant proportion of anucleate cells (2.4% of the population excluding lysed cells), whereas only a mild over-replication phenotype was observed in the presence of xylose with fewer cells lacking DNA (Fig. 2E-F). Together with cell growth analyses (Fig. 1C-D), the results indicate that membrane depolarisation occurs after DNA damage is detected, at a similar time as the onset of cell lysis.

### The lethal phenotype caused by DnaA^G154S^ is independent of prophage induction

The bacterial DNA damage response is known to activate lysogenic bacteriophages that can contribute to cell lysis. To determine whether the lysis phenotype observed in the *Bs^HI^* strain was due to prophage activation, the hyper-initiation system was transformed into a *B. subtilis* strain lacking the six prophage elements harboured in the parental lab strain (Δ6) [43, 44]. Spot titre analyses showed that after eight hours of growth in the absence of inducer, the *Bs^HI^* Δ*6* strain displayed a ∼100-fold decrease in CFU (Fig. 3A). MFA validated that *Bs^HI^* Δ*6* cells experienced significant hyper-initiation (Fig. 3B). These results indicate that prophage induction cannot explain the growth defects observed in cells hyper-initiating DNA replication. Nonetheless, to avoid issues of prophage induction, all subsequent experiments were performed using the Δ6 genetic background.

**FIGURE 3:**
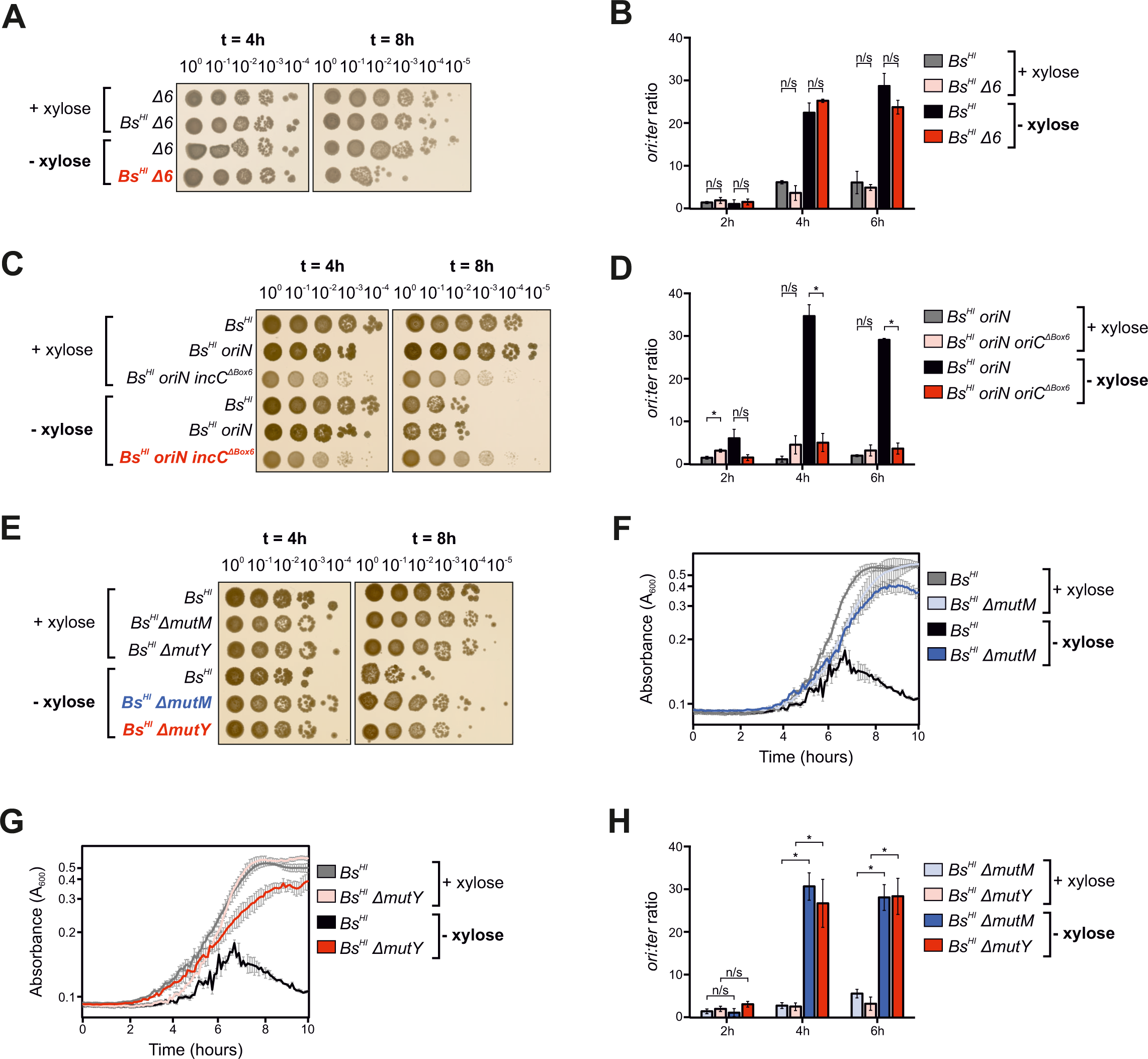
Cell lysis caused by hyper-initiation is not due to prophage induction, is dependent on the replication origin *oriC* and can be rescued by limiting DNA repair. (A, C, E) Spot titre analyses showing that the hyper-initiation induced lethal phenotype **(A)** is not due to the activity of major prophages, **(C)** is dependent on the presence of the functional replication origin oriC recognised by DnaA, **(E)** can be rescued by a knockout of DNA repair genes *mutM* or *mutY*. **(B, D, H)** Marker frequency analyses showing that **(B)** a strain lacking major prophages remains capable of hyper-initiating DNA replication, **(D)** the presence of the ectopic origin *oriN* does not affect the ability of *Bs^HI^*cells to hyper-initiate, whereas combination with an inactive endogenous origin (*oriN oriC*^Δ*Box6*^) abolishes hyper-initiation, **(H)** deletion of *mutM* or *mutY* does not affect the ability of *dnaA^G154S^ P_xyl_-parA^G12V^* cells to hyper-initiate. **(F-G)** Growth curves showing that the deletion of (F) *mutM* or (G) *mutY* is associated with a fitness cost slowing the growth of *Bs^HI^*cells in the absence of xylose. (A-B) Strains: Δ6 (CW483), *Bs^HI^* Δ6 (HM1971). (C-D) Strains: *Bs^HI^* (HM1970), *Bs^HI^ oriN* (HM2014), *Bs^HI^ oriN oriC*^Δ*box6*^ (HM2015). (E-H) Strains: *Bs^HI^* (HM1971), *Bs^HI^* Δ*mutM* (HM2012), *Bs^HI^* Δ*mutY* (HM2011). Non-significant (n/s), P values < 0.05 (*).

### The lethal phenotype caused by DnaA^G154S^ requires DNA replication hyper-initiation from *oriC*

In addition to being the master initiator of DNA replication in bacteria, DnaA also plays roles in diverse cellular processes such transcription regulation, chromosome organisation, cell division, cell differentiation, and metabolism [45, 46]. To ascertain whether the lethal phenotype observed with DnaA^G154S^ was dependent upon hyper-initiation from the endogenous replication origin *oriC*, a strain that replicates from a DnaA-independent origin *oriN* (*Bs^HI^ oriN*) was constructed, thus allowing mutagenesis of an essential DnaA-box within *oriC* (*Bs^HI^ oriN incC*^Δ*Box6*^) [47]. Grown in the absence of xylose for eight hours, the *Bs^HI^ oriN* strain displayed a similar drop in CFU to the parental strain, indicating that the presence of *oriN* does not alleviate the hyper-initiation phenotype (Fig. 3C). In contrast, the *Bs^HI^ oriN incC*^Δ*Box6*^ strain showed no decrease in CFU following growth in the absence of xylose (Fig. 3C). MFA confirmed that the *Bs^HI^ oriN* strain hyper-initiates DNA replication, whereas the addition of the *incC*^Δ*Box6*^ mutation alleviated the ability of DnaA^G154S^ to over activate *oriC* (Fig. 3D). These results indicate that hyper-initiation of DNA replication from *oriC* is necessary for DnaA^G154S^ to elicit cell death.

### Suppression of lethal DNA replication hyper-initiation by limiting base excision repair

Studies in *E. coli* have suggested that DNA replication hyper-initiation causes DNA damage when the replisome encounters DNA repair events occurring on the chromosome [17, 18]. Under aerobic growth conditions, nucleobase oxidation has been proposed to be a source of DNA damage requiring repair [48, 49]. To initiate repair of damaged DNA, well-studied enzymes including the MutM and MutY glycosylases must first excise damaged bases before replacing them, potentially resulting in single-stranded DNA discontinuities that can promote replisome collapse and emergence of DNA double-strand breaks (DSB), which elicit recruitment of further critical repair complexes [50–52]. To explore this model, genes encoding base excision repair factors MutM or MutY were deleted in the *Bs^HI^* strain and whole genome sequencing validated the absence of suppressor mutations in essential genes (Fig. S6A-B, *mutM*/*mutY* mutants are known to be mutagenic [53]). Spot titre analyses revealed that removing either of these DNA repair genes partially supressed the lethal phenotype observed in the parental *Bs^HI^* strain (Fig. 3E, note the 5 to 10 fold reduction in CFU compared to cells grown with xylose) and plate reader analyses identified that these mutations had a fitness cost resulting in slower growth (Fig. 3F-G). Critically, MFA showed that the strains harbouring either *mutM* or *mutY* knockouts remained capable of hyper-initiating DNA replication (Fig. 3H). Taken together and consistent with observations made in *E. coli* [17], the data suggest that excessive replication from *oriC* can lead to conflicts between DNA replication and base excision repair. Here in *B. subtilis*, the results show general elevation of the bacterial stress response by production of DNA damage substrate of RecA, induction of the SOS response, accumulation of morphological changes and ultimately cell death via lysis.

### Hyper-initiation induced cell death is conserved in *S. aureus* but independent of lysis

The replication initiator protein DnaA is broadly conserved in prokaryotes and we wondered whether DNA hyper-initiation could be exploited to inhibit the growth of bacterial pathogens. To test this idea, the consequence of DNA replication hyper-initiation was determined in *S. aureus*. An ectopic copy of *S. aureus* wild-type *saDnaA* (encoding SaDnaA) or *saDnaA^G159S^*(encoding SaDnaA^G159S^, analogous to the hypermorphic *B. subtilis* DnaA^G154S^ variant) was placed under the control of an anhydrotetracycline-inducible (aTc) promoter on a self-replicating plasmid (Fig. 4A), and the empty inducible plasmid was used as control to recapitulate conditions where only the native endogenous *saDnaA* is present [26, 54].

**FIGURE 4:**
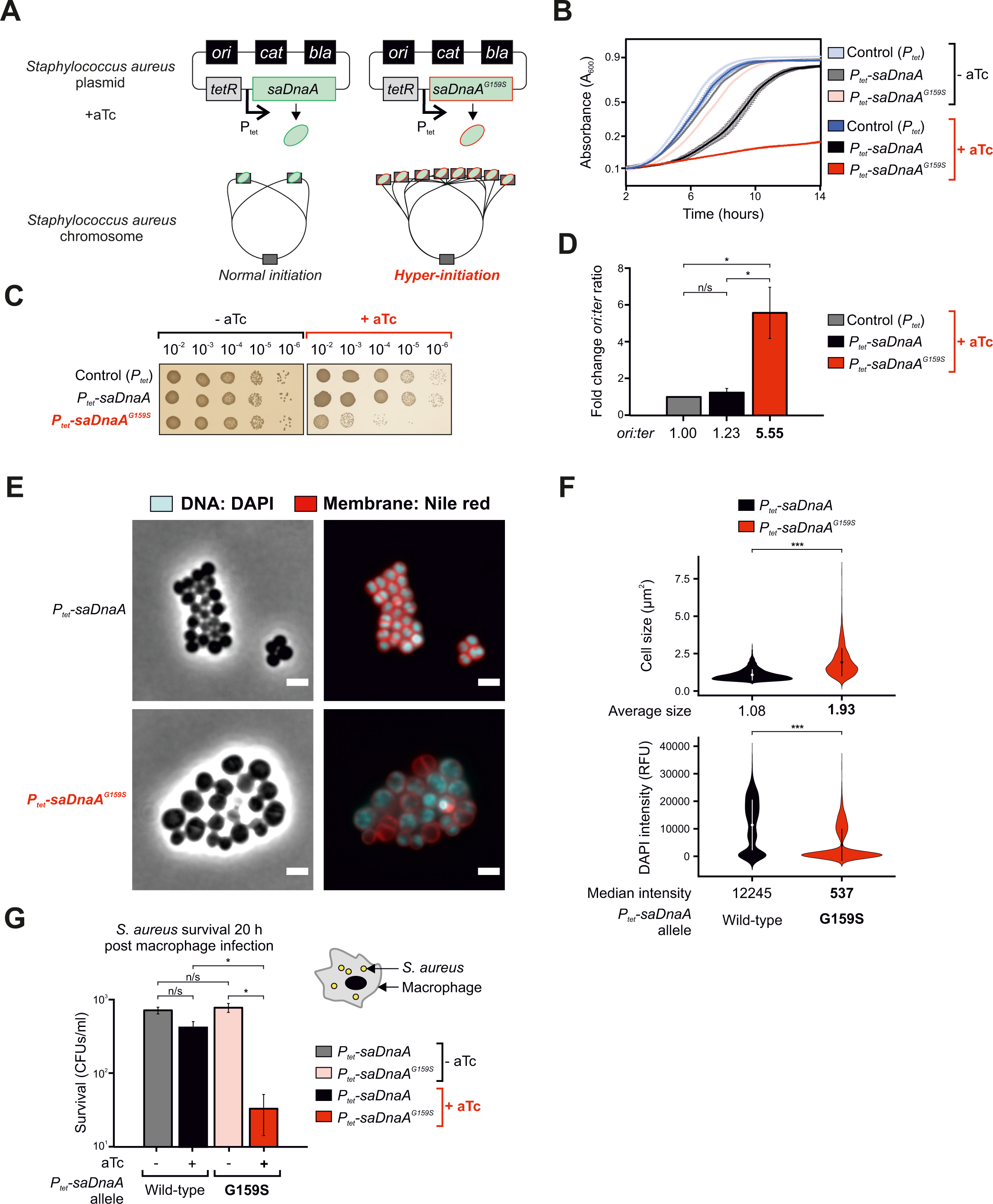
Hyper-initiation in *S. aureus* causes cell death and reduces survival in murine macrophages. **(A)** Genetic system used to artificially modulate DNA replication hyper-initiation in *S. aureus*. Anhydrotetracycline (aTc) induction controls expression of *saDnaA* variants (wild-type or hyper-initiator allele *saDnaA^G159S^*). **(B)** Plate reader analyses showing the absence of cell growth in cells harbouring the hyper-initiation allele *saDnaA^G159S^* in the presence of aTc. **(C)** Spot titre assay showing that the hyper-initiation allele *saDnaA^G159S^* inhibits the growth of *S. aureus*. **(D)** Marker frequency analyses showing that *saDnaA^G159S^* cells hyper-initiate DNA replication in the presence of aTc. Fold change ori:ter ratios are normalised to the empty vector control (*P_tet_*). **(E)** Microscopy experiments showing that hyper-initiation (*P_tet_-saDnaA^G159S^* cells) leads to the accumulation of morphological changes. Fluorescence images are merged between cyan channel for visualisation of DNA content (DAPI dye) and red signal for membrane stain (Nile red). Scale bar indicates 2 μm. **(F)** Microscopy analyses showing quantification of individual cell sizes (top) and UV fluorescence (bottom, nucleoid signal via DAPI) from the data shown in (E). Each condition highlights the mean (circles within violins) and standard deviation (vertical line crossing the mean). **(G)** Survival plot showing that *S. aureus* infection is attenuated when inducing the hyper-initiator allele *saDnaA^G159S^*. (B-F) Strains: Control *P_tet_* (CW1095), *P_tet_-saDnaA* (CW1150), *P_tet_-saDnaA^G159S^* (CW1151). Non-significant (n/s), P values: < 0.05 (*), < 0.0001 (***).

To characterise the effect of the SaDnaA^G159S^ variant, strains were grown in liquid media in the absence or presence of aTc (e.g. repression or induction of *saDnaA*/*saDnaA^G159S^*). Plate reader analyses showed that *S. aureus* harbouring the empty vector control was minimally affected by this plasmid system in the absence or the presence of aTc, with comparable growth dynamics to cells encoding the uninduced *saDnaA* construct (Fig. 4B). Expression of SaDnaA^G159S^ significantly inhibited growth of the culture, while expression of wild-type SaDnaA led to a longer lag phase but comparable doubling-time during exponential growth (Fig. 4B). These results are consistent with a previous report that overexpression of SaDnaA in *S. aureus* is toxic [54].

To quantify the effect of SaDnaA^G159S^ on cell viability, cultures were grown overnight in the absence of aTc, then serially diluted and aliquoted onto solid media with or without inducer. Spot titre analyses showed that in the absence of inducer to express *saDnaA* alleles, each strain produced a similar number of CFU, indicating that these plasmids are not cytotoxic under the conditions tested (Fig. 4C). In the presence of aTc, both the strain with the empty vector and the strain expressing wild-type SaDnaA showed a small colony phenotype, but nonetheless produced similar CFU compared to cultures grown without inducer. In contrast, expression of SaDnaA^G159S^ was associated with a ∼100-fold loss in CFU (Fig. 4C, note that this loss of viability is comparable to the growth inhibition observed using the *B. subtilis* hyper-initiation system in Figs. 1D, 3A). Consistent with the literature yet contrasting with the Gram-negative bacterium *E. coli*, overexpression of wild-type *saDnaA* did not significantly affect replication initiation in *S. aureus* (Fig. 4D) [18, 55]. MFA demonstrated that induction of the hyper-active allele *saDnaA^G159S^* led to a significant ∼5 fold increase in levels of DNA replication initiation compared to controls, whereas induction of wild-type *saDnaA* yielded similar results to the empty vector condition (Fig. 4D). Together, these data indicate that DNA replication hyper-initiation inhibits bacterial growth in *S. aureus*.

To follow the consequence of hyper-initiation at a single cell level, timelapse microscopy was used to observe the growth of a strain harbouring the plasmid expressing SaDnaA^G159S^. Cultures were grown to early exponential phase, then spotted onto agarose pads in the presence or absence of inducer aTc. Over eight hours, uninduced *S. aureus* cells were able to grow and fill in the field of view without notable morphological abnormalities (Supplementary Video 3). However in the presence of aTc, strong growth inhibition and a significant increase in cell size were observed (Supplementary Video 4). Note that in this context cell lysis was not detected, likely reflecting inherent differences between the cell envelopes of *S. aureus* and *B. subtilis* (i.e. thick staphylococcal cell wall).

To further understand the growth inhibition observed during hyper-initiation in *S. aureus*, fluorescence microscopy was employed to visualise DNA content and the cell membrane using the dyes DAPI and Nile red, respectively. Wild-type and hyper-initiation SaDnaA proteins were induced with aTc in early exponential phase and cells were imaged after 90 minutes. Under these conditions, cells expressing wild-type SaDnaA showed uniform DAPI staining (i.e. normal chromosome content, Fig. 4E). In contrast, cells expressing the SaDnaA^G159S^ variant appeared significantly larger (approximately twice the average cell size of the wild-type control) with heterogeneous DNA content including many cells devoid of DNA, many of which featuring extremely low DAPI intensity (Fig. 4E-F, median fluorescence intensity over 22 times lower than control). Thus, DNA replication hyper-initiation in *S. aureus* leads to chromosome loss and cell death.

### Hyper-initiation decreases *S. aureus* survival in murine macrophages

*S. aureus* is an opportunistic pathogen able to survive within immune cells and can cause respiratory, gut, blood and skin infections [56]. During the immune response, macrophages engulf *S. aureus* and are thought to release reactive oxygen species, potentially creating DNA damage. We hypothesised that DNA replication hyper-initiation in *S. aureus* might provide synergy with DNA damage generated in macrophages to inhibit bacterial survival during infection.

To test this model, RAW 264.7 murine macrophage-like cells were infected with *S. aureus* harbouring plasmids encoding either wild-type or *saDnaA^G159S^* alleles. Following a one-hour incubation to enable phagocytosis, extracellular bacteria were killed using gentamycin treatment and cultures were incubated for 20 hours in the presence or absence of aTc. Macrophages were then harvested and lysed to assess the number of surviving intracellular *S. aureus* cells. In uninduced conditions, strains expressing SaDnaA and SaDnaA^G159S^ yielded similar *S. aureus* survival (Fig. 4G). In the presence of aTc, macrophages infected with wild-type *saDnaA* showed a negligeable reduction in bacterial load compared to uninduced cells (1.4x less CFUs). In contrast, expression of SaDnaA^G159S^ produced a 25-fold reduction in *S. aureus* survival compared to the control (Fig. 4G). Together, these results suggest that *S. aureus* experiences oxidative DNA damage within macrophages which is exacerbated by DNA replication hyper-initiation, thereby inhibiting bacterial proliferation.

## DISCUSSION

Here we report that hyper-initiation of DNA replication, achieved via expressing variants of the master initiator DnaA, inhibits proliferation of both *B. subtilis* and *S. aureus*. The data support previous hyper-initiation studies focused on the Gram-negative bacterium *E. coli* [17, 18] and add incremental knowledge to a recent investigation that identified pathways able to rescue over-initiation in *B. subtilis* [19]. Interestingly, we identify that the pathways underlying hyper-initiation induced cell death across Gram-positive species share some similarities but appear to be distinct.

In *B. subtilis*, hyper-initiation dysregulates several important cellular processes including broad elevation of the DNA damage response, accumulation of morphological deformations and membrane depolarisation, which together lead to cell death via lysis (Figs. 1-2). Importantly, these mechanisms cannot be attributed to the activation of prophage and are *oriC*-dependent, indicating that DNA replication hyper-initiation is causative of the induced lethality (Fig. 3A-D). Consistent with observations made in *E. coli* and mycobacteria [17, 18, 57], limiting base excision repair partially suppresses growth inhibition, suggesting that the repair activity of MutM/MutY generates a second-line challenge to oncoming replication forks by promoting fork collapse (Fig. 3E-H). These results imply that a source of stress during hyper-initiation likely results from conflicts between the replication and repair machineries, leading to the emergence of DSBs that elicit critical RecA-mediated homologous repair processes (Fig. S4).

Inspired by these findings, we further found that hyper-initiation can be exploited to limit proliferation of the human pathogen *S. aureus* (Fig. 4). The ability of *S. aureus* to survive within immune cells is thought to contribute to bacterial persistence [56]. Therefore, exploring approaches to reduce the intracellular pool of bacteria could favour clearance of *S. aureus* associated infections. Under standard laboratory growth conditions, we note that hyper-initiation of DNA replication generates significant morphological defects associated with cell death. During infection, experiments showed that hyper-initiation can also decrease *S. aureus* survival in macrophages. We speculate that hyper-initiation and associated replication/repair conflicts provides synergy with reactive oxygen species produced by macrophages during infection, thereby resulting in increased DNA damage and strong bacterial growth inhibition.

Our findings open new avenues for the development of alternative antimicrobial strategies. In this context, while several adverse approaches have been proposed to inhibit bacterial DNA replication [58], hyper-activation of DNA replication remains relatively unexplored. Interestingly, clinical isolates of *Mycobacterium tuberculosis* with mutations in *dnaA* were found to modulate resistance upon exposure to different classes of antimicrobials [59]. Therefore, the development of novel compounds targeting DnaA to exploit hyper-initiation as an antibiotic adjuvant may be an attractive alternative to potentiate antimicrobials targeting DNA synthesis.

## AUTHOR CONTRIBUTIONS

HG, DS, JH, KM, HM and CW contributed to the conception/design of the work. HG, DS, JG, KM, HM and CW generated results presented in the manuscript. HG and CW created figures. HG, HM and CW wrote the manuscript. JG, KM, HM and CW edited the manuscript.

## ACKNOWLEDGEMENTS

The authors thank Frances Davison and Amandeep Kaur for technical assistance.

## DATA SUMMARY

Supplementary Videos are available on Figshare [60].

## SUPPLEMENTARY VIDEOS

**Supplementary Video 1.** Hyper-initiation in a *B. subtilis* strain growing in presence of xylose (repressed hyper-initiation). Strain: HG282.

**Supplementary Video 2.** Hyper-initiation in a *B. subtilis* strain growing in absence of xylose (induced hyper-initiation). Strain: HG282.

**Supplementary Video 3.** Hyper-initiation in a *S. aureus* strain growing in absence of aTc (repressed hyper-initiation). Strain: CW1151.

**Supplementary Video 4.** Hyper-initiation in a *S. aureus* strain growing in presence of aTc (induced hyper-initiation). Strain: CW1151.

---

**SUPPLEMENTARY FIGURE 1:**
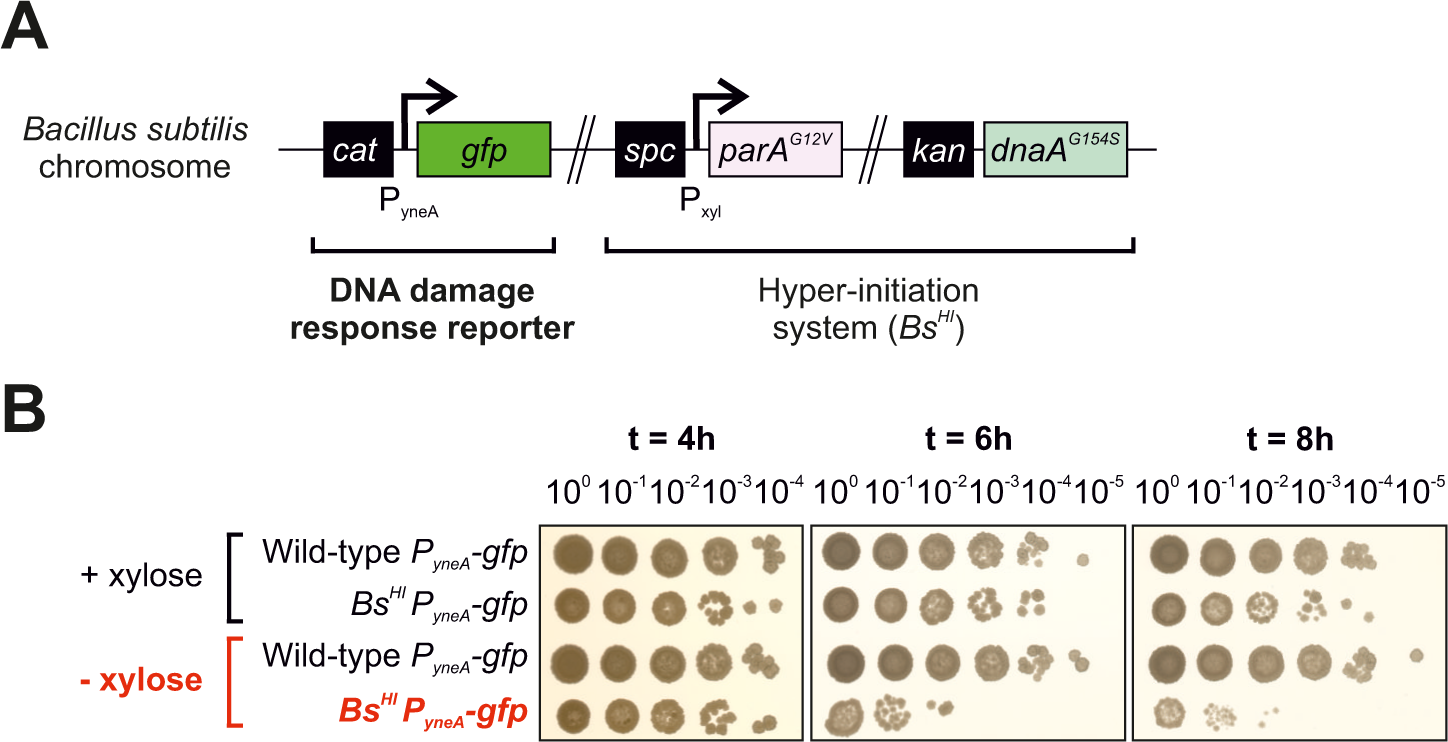
DNA damage response reporter in *B. subtilis*. **(A)** Genetic system used to artificially modulate hyper-initiation and to visualise the onset of the DNA damage response simultaneously via the *P_yneA_-gfp* reporter cassette. **(B)** Spot titre assay showing that the *P_yneA_-gfp* reporter cassette does not affect the lethal phenotype observed upon induction of hyper-initiation using xylose. Strains: Wild-type *P_yneA_-gfp* (HG53), *Bs^HI^ P_yneA_-gfp* (HM1981).

**SUPPLEMENTARY FIGURE 2:**
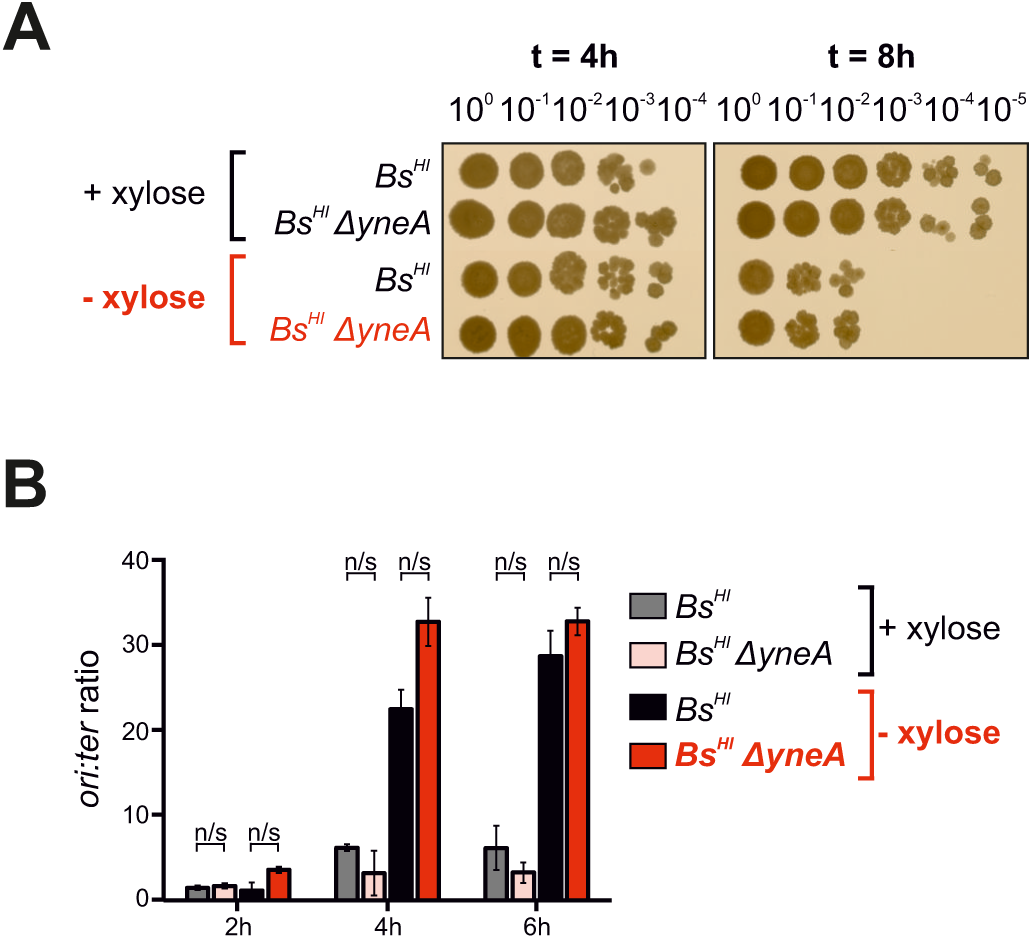
YneA is not causative of hyper-initiation induced cell death. **(A)** Spot titre assay showing that the phenotype associated with *Bs^HI^* cells remains unchanged upon the knockout of the DNA damage response factor *yneA* in the presence/absence of xylose to control hyper-initiation. **(B)** Marker frequency analyses demonstrate that the absence of *yneA* does not affect hyper-initiation controlled via xylose induction in *Bs^HI^* cells (non-significant: n/s). Strains: *Bs^HI^* (HM946), *Bs^HI^* Δ*yneA* (HG424).

**SUPPLEMENTARY FIGURE 3:**
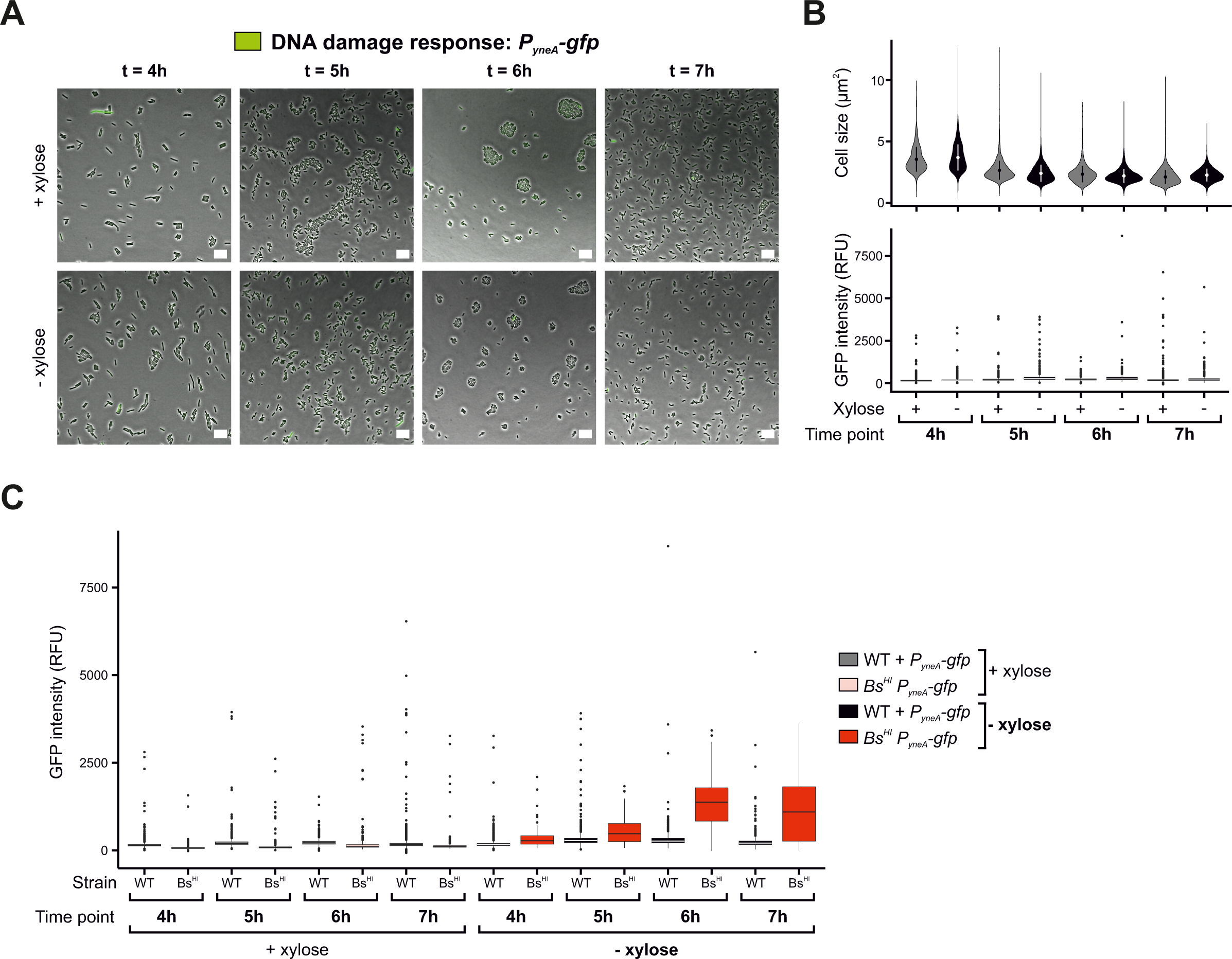
DNA damage response reporter in wild-type *B. subtilis* and comparison with the hyper-initiation strain. **(A)** Microscopy experiments showing that the DNA damage response reporter *P_yneA_-gfp* produces a low background signal in wild-type *B. subtilis* in the presence/absence of xylose. Phase contrast images are overlayed with a green fluorescence channel corresponding to the *P_yneA_-gfp* signal. Scale bar indicates 10 μm. Strain: HM1964. **(B)** Microscopy analyses showing quantification of individual cell sizes (top) and GFP fluorescence (bottom, DNA damage reporter signal) from the data shown in (A). **(C)** Microscopy analyses showing quantification of GFP fluorescence from the *P_yneA_-gfp* reporter cassette for data extracted from wild-type (WT) or hyper-initiation (Bs^HI^) strains. Strains: Wild-type (WT) + *P_yneA_-gfp* (HM1964), *Bs^HI^ P_yneA_-gfp* (HG287). (B/C) Each condition highlights mean cell sizes (circles within violins), standard deviation (vertical line crossing the mean) and boxplots indicate median lines (within boxes) and outliers (circles outside boxes).

**SUPPLEMENTARY FIGURE 4:**
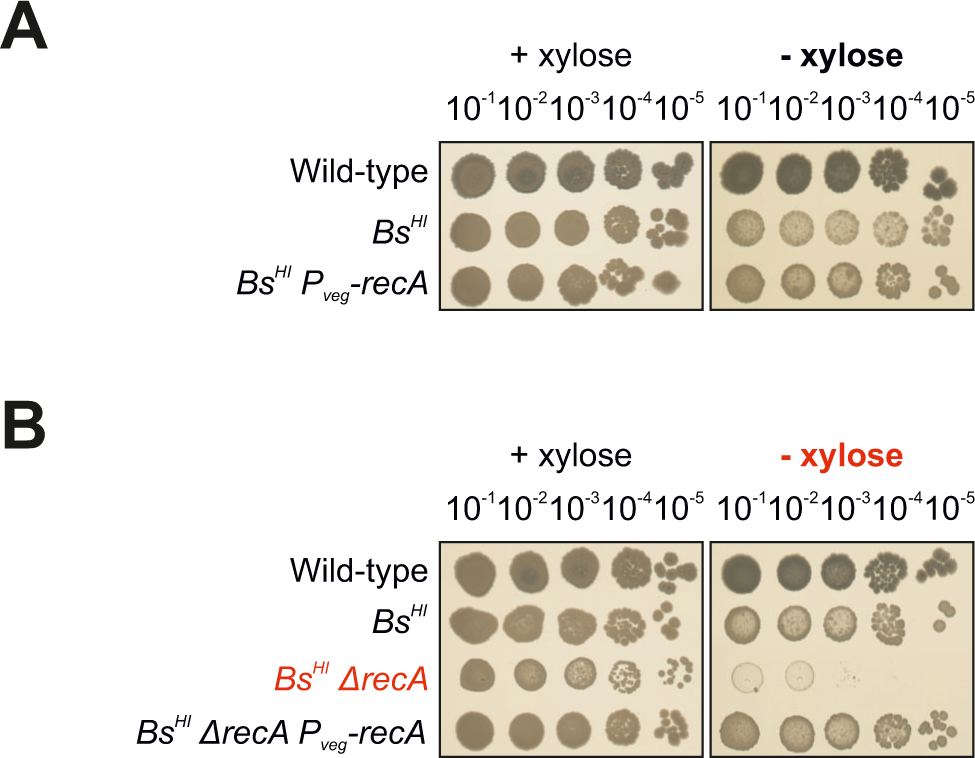
Deletion of *recA* leads to synthetic lethality in the *B. subtilis* hyper-initiation strain. **(A)** Spot titre assay showing that addition of an ectopic copy of *recA* does not affect viability of wild-type or *Bs^HI^* cells in the presence/absence of xylose to control hyper-initiation. **(B)** Spot titre assay showing that *recA* is essential for viability in *Bs^HI^*cells in the absence of xylose during hyper-initiation. Strains: Wild-type (CW483), *Bs^HI^* (HM1970), *Bs^HI^ P_veg_-recA* (CW2160), *Bs^HI^* Δ*recA* (CW2156), *Bs^HI^* Δ*recA P_veg_-recA* (CW2163).

**SUPPLEMENTARY FIGURE 5:**
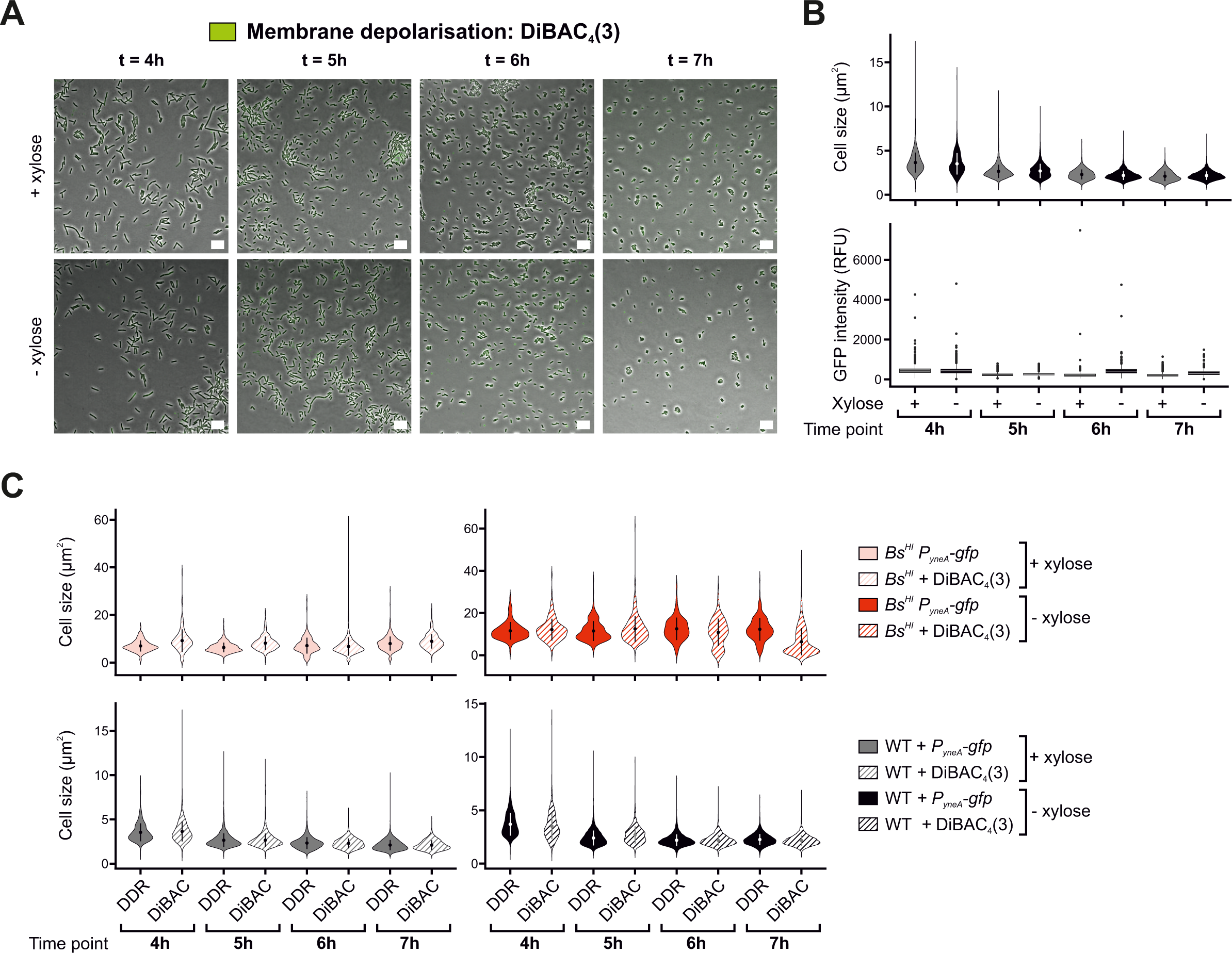
Membrane depolarisation in wild-type *B. subtilis* and comparison of cell sizes across time course assays. **(A)** Microscopy experiments showing that the membrane depolarisation dye DiBAC_4_(3) produces a low background signal in wild-type *B. subtilis* in the presence/absence of xylose. Phase contrast images are overlayed with a green fluorescence channel corresponding to the DiBAC_4_(3) signal. Scale bar indicates 10 μm. Strain: HM715. **(B)** Microscopy analyses showing quantification of individual cell sizes (top) and GFP fluorescence (bottom, membrane depolarisation signal) from the data shown in (A). **(C)** Microscopy analyses comparing cell sizes between hyper-initiation (top) and wild-type strains (bottom) in the presence/absence of xylose from both DNA damage response (DDR) and membrane depolarisation experiments (DIBAC). Strains: *Bs^HI^ P_yneA_-gfp* (HG287), *Bs^HI^* + DiBAC_4_(3) (HM946), Wild-type (WT) + *P_yneA_-gfp* (HM1964), WT + DiBAC_4_(3) (HM715). (B/C) Each condition highlights mean cell sizes (circles within violins), standard deviation (vertical line crossing the mean) and boxplots indicate median lines (within boxes) and outliers (circles outside boxes).

**SUPPLEMENTARY FIGURE 6:**
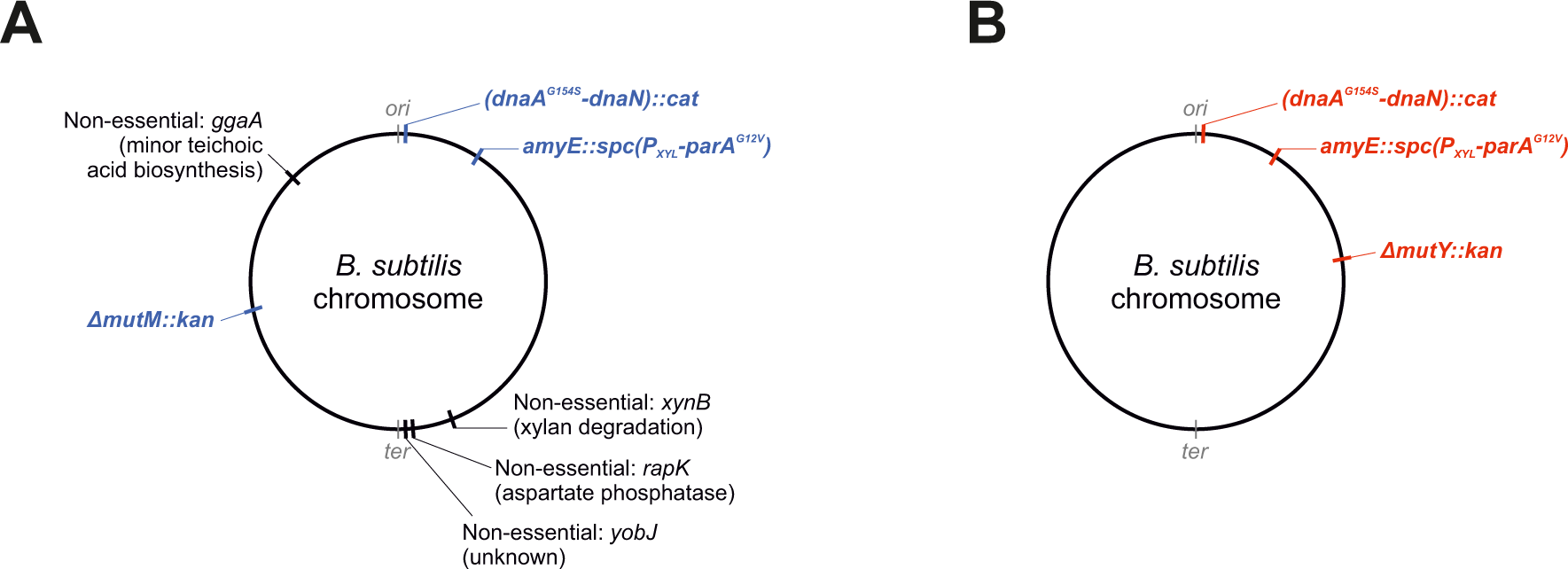
Validation of *mutM* and *mutY* knockouts via whole genome sequencing. **(A-B)** Mutations detected via whole genome sequencing performed on hyper-initiation strains *Bs^HI^*Δ*mutM* (A) and *Bs^HI^* Δ*mutY* (B). Intended genetic modifications are indicated in blue and red for Δ*mutM* and Δ*mutY* backgrounds, respectively. Background mutations were only found in the *mutM* mutant (black annotations of mutated genes alongside predicted/known functions). Strains: *Bs^HI^* Δ*mutM* (HM2012), *Bs^HI^* Δ*mutY* (HM2011).

**Supplementary Table 1.**
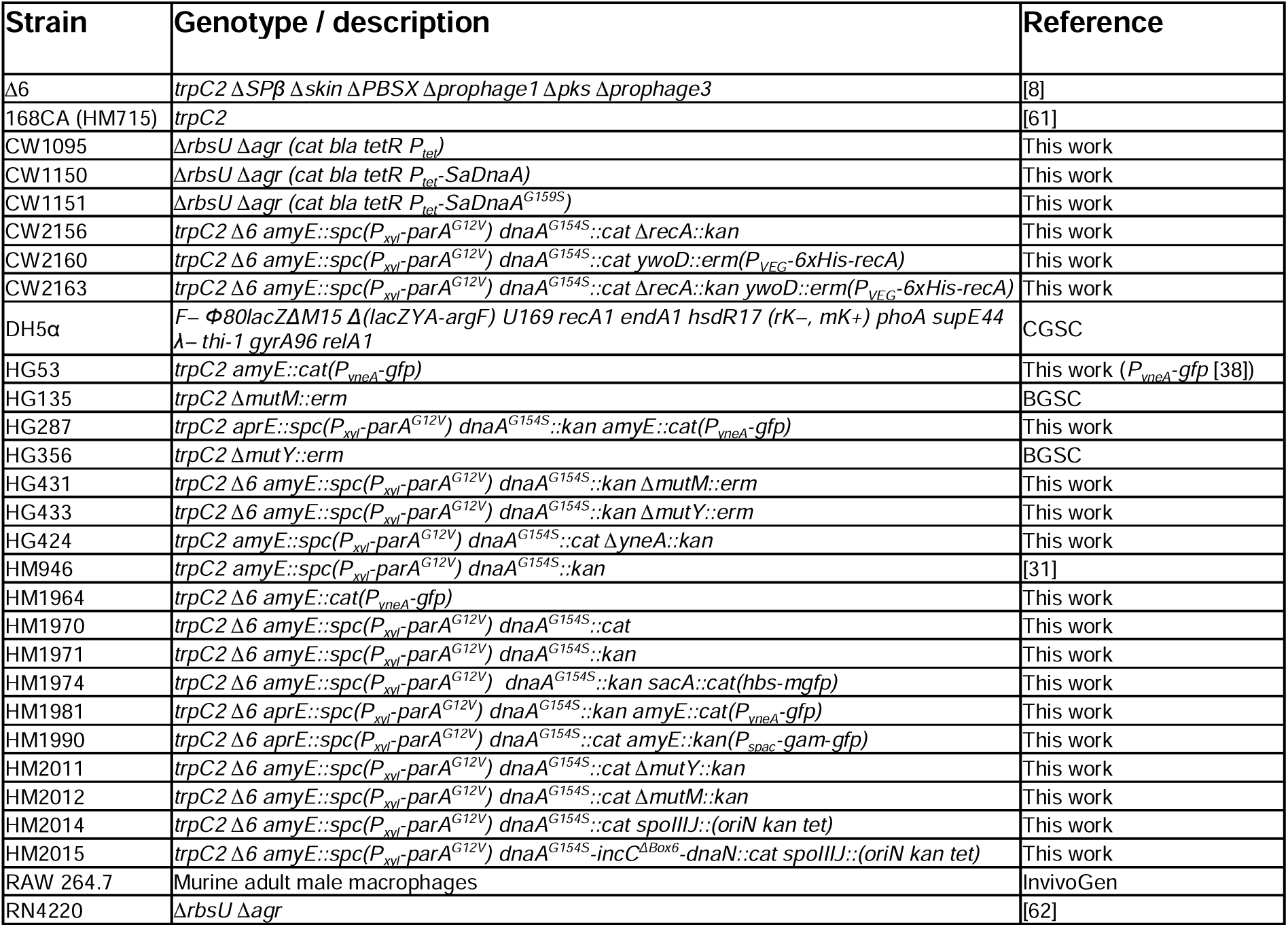
Strains used in this study.

**Supplementary Table 2.**
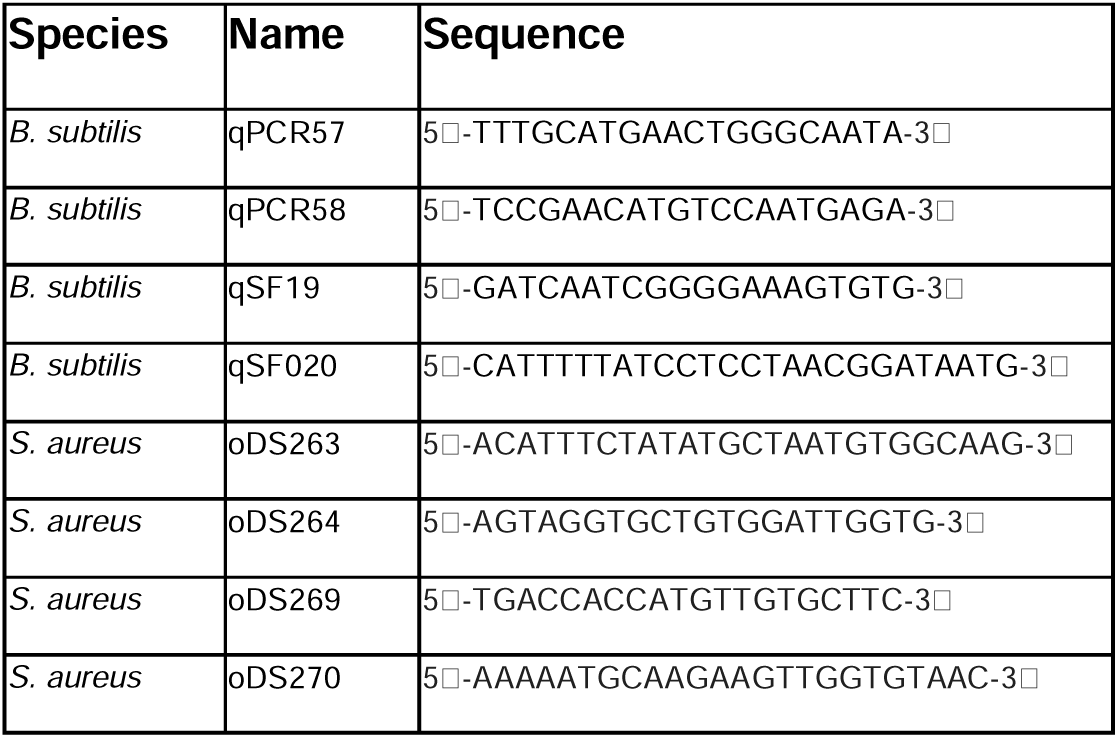
Primers used for qPCR.

